# Live-dead assay on unlabeled cells using phase imaging with computational specificity

**DOI:** 10.1101/2020.10.28.359554

**Authors:** Chenfei Hu, Shenghua He, Young Jae Lee, Yuchen He, Edward M. Kong, Hua Li, Mark A. Anastasio, Gabriel Popescu

**Affiliations:** Department of Electrical and Computer Engineering, Washington University in St. Louis, St. Louis, Missouri, 63130, USA; Beckman Institute for Advanced Science and Technology, Washington University in St. Louis, St. Louis, Missouri, 63130, USA; Department of Computer Science & Engineering, Washington University in St. Louis, St. Louis, Missouri, 63130, USA; Neuroscience Program, University of Illinois at Urbana-Champaign, Urbana, Illinois, 61801, USA; Department of Bioengineering, University of Illinois at Urbana-Champaign, Urbana, Illinois, 61801, USA; Carle Cancer Center, Carle Foundation Hospital, Urbana, IL, 61801, USA

## Abstract

Existing approaches to evaluate cell viability involve cell staining with chemical reagents. However, this step of exogenous staining makes these methods undesirable for rapid, nondestructive and long-term investigation. Here, we present instantaneous viability assessment of *unlabeled* cells using phase imaging with computation specificity (PICS). This new concept utilizes deep learning techniques to compute viability markers associated with the specimen measured by label-free quantitative phase imaging. Demonstrated on different live cell cultures, the proposed method reports approximately 95% accuracy in identifying live and dead cells. The evolution of the cell dry mass and projected area for the labelled and unlabeled populations reveal that the viability reagents decrease viability. The nondestructive approach presented here may find a broad range of applications, from monitoring the production of biopharmaceuticals, to assessing the effectiveness of cancer treatments.

## Introduction

Rapid and accurate estimation of viability of biological cells is important for assessing the impact of drug, physical or chemical stimulants, and other potential factors on cell dynamics. The existing methods to evaluate cell viability commonly require mixing a population of cells with reagents to convert a substrate to a colored or fluorescent product [1]. For instance, using membrane integrity as an indicator, the live and dead cells can be separated by trypan blue exclusion assay, where only nonviable cells are stained and appear as a distinctive blue color under a microscope [2, 3]. Starting in 1970s, fluorescence imaging has developed as a more accurate, faster, and reliable method to determine cell viability [4-7]. Similar to the principle of trypan blue test, this method identifies nonviable cells by using fluorescent reagents only taken up by cells that lost their membrane permeability barrier. Unfortunately, the step of exogenous staining generally requires some incubation time for optimal staining intensity, making these methods difficult for quick evaluation. More importantly, the toxicity introduced by stains eventually kills the cells and, thus, prevents the long term investigation.

Quantitative phase imaging (QPI) is a label-free modality that has gained significant interest due to its broad range of potential biomedical applications [8, 9]. QPI measures the optical phase delay across the specimen as an intrinsic contrast mechanism, and thus, allows visualizing transparent specimen (i.e., cells and thin tissue slices) with nanoscale sensitivity, which makes this modality particularly useful for nondestructive investigations of cell dynamics (i.e. growth, proliferation, and mass transport) in both 2D and 3D [10-15]. In addition, the optical phase delay is linearly related to the non-aqueous content in cells (referred to as dry mass), which directly yields biophysical properties of the sample of interest [16-18]. More recently, with the concomitant advances in deep learning, we have witnessed exciting new avenues for QPI. As a QPI map encodes structure and biophysical information, it is possible to apply deep learning techniques to extract subcellular structures, perform signal reconstruction, correct image artifacts, convert QPI data into virtually stained or fluorescent images, and diagnose and classify various specimens [19-27].

In this article, we demonstrate that rapid viability assay can be conducted in a label-free manner using spatial light interference microscopy (SLIM) [28, 29], a highly sensitive QPI method, and deep learning. We apply the concept of our newly-developed phase imaging with computational specificity (PICS) to digitally stain for the live and dead markers. Demonstrated on live adherent HeLa and CHO cell culture, we predict the viability of individual cell measured with SLIM by using a joint EfficientNet [30] and transfer learning [31] strategy. Using the standard fluorescent viability imaging as ground truth, the trained neural network classifies the viable state of individual cell with 95% accuracy. Furthermore, by tracking the cell morphology over time, unstained HeLa cells show significantly higher viability compared to the cells stained with viability reagents. These findings suggest that PICS method enables rapid, nondestructive, and unbiased cell viability assessment, potentially, of great benefit to the general biomedical community.

## Results

The procedure of image acquisition is summarized in Fig. 1. We employed spatial light interference microscopy (SLIM) [28] to measure quantitative phase map of cells *in vitro*. The system is built by attaching a SLIM module (CellVista SLIM Pro, Phi Optics, Inc.) to the output port of an existing phase-contrast microscope (Fig. 1a). By modulating the optical phase delay between the incident and the scattered field, a quantitative phase map is retrieved from four intensity images via phase-shifting interferometry [32]. SLIM employs a broadband LED as illumination source and common-path imaging architecture, which yields sub-nanometer sensitivity to optical pathlength change and high temporal stability [32, 33]. By switching to epi-illumination, the optical path of SLIM is also used to record the fluorescent signals over the same field of view. Detailed information about the microscope configuration can be found in *Methods*.

**Figure 1.**
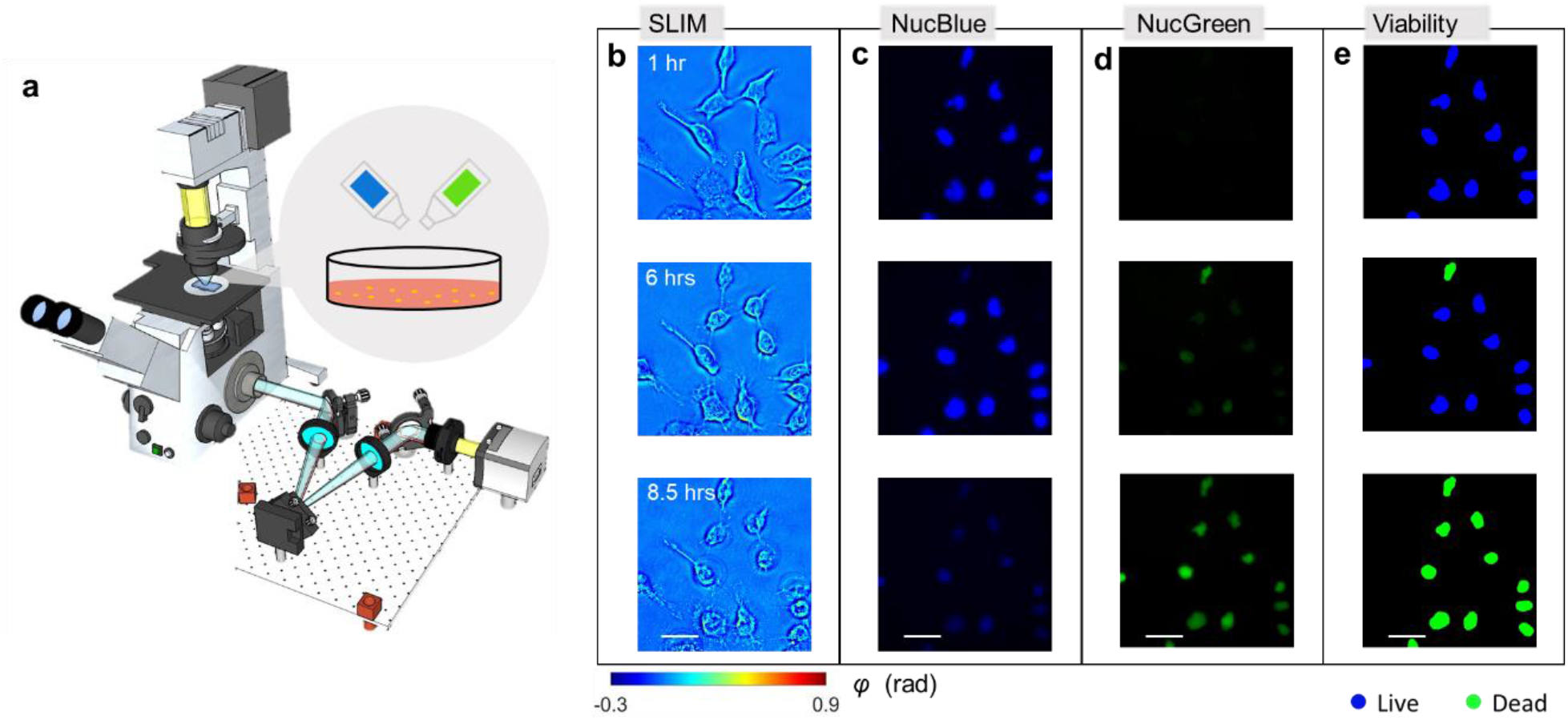
Schematic of the imaging system and representative results. **a**. CellVista SLIM Pro microscope (Phi Optics, Inc.) consists of an existing phase contrast microscope and an external module attached to the output port. By switching between transmission and reflection excitation, both SLIM and co-localized fluorescence images can be recorded via the same optical path. Before time-lapse imaging started, fluorescence viability reagents were mixed with HeLa cell culture. **b**. Representative SLIM measurements of HeLa cell at 1, 6, and 8.5 hours. **c**. NucBlue fluorescent signals of the live viability reagent. **d**. NucGreen fluorescent signals of the dead viability reagents measured by a FITC filter. **e**. Viability states of the individual cells. Scale bars: 50 microns

To demonstrate the feasibility of the proposed method, we imaged and analyzed live cell cultures. Before imaging, two drops of cell-viability-assay reagents (ReadyProbes Cells Viability Imaging Kit, Thermofisher) were added into the growth media, and the cells were then incubated for approximately 15 minutes to achieve optimal staining intensity. The viability-assay kit contains two fluorescently labeled reagents: NucBlue (the “live” reagent) combines with the nuclei of all cells and can be imaged with a DAPI fluorescent filter set, and NucGreen (the “dead” reagent) stains the nuclei of cells with compromised membrane integrity, which is imaged with a FITC filter set. In this assay, live cells produce blue fluorescent signal; dead cells emit both green and blue fluorescence; The procedure of cell culture preparation can be found in *Methods*.

After staining, the sample was transferred to the microscope stage, and measured by SLIM and epi-fluorescence microscopy. In order to generate a heterogeneous cell distribution that shifts from predominantly alive to mostly dead cells, the imaging was performed under room conditions, such that the low-temperature and imbalanced pH level in the media would adversely injure the cells and eventually cause necrosis. Recording one measurement every 30 or 60 minutes, the entire imaging process lasted for approximately 10 hours. We repeated this experiment for four times to capture the variability among different batches. Figure 1b shows the SLIM images of HeLa cells measured at t = 1 hour, 6, and 8.5 hours, respectively, and the corresponding fluorescent measurements are shown in Figs. 1c-d. The results in Fig. 1 show that the adverse environmental condition continues injuring the cell, where blebbing and membrane disruption could be observed during cell death. Our QPI measurements agree with the results reported in previous literature [34]. On the other hand, these morphological alterations are correlated with the changes in fluorescence signals, where the intensity of NucGreen (“dead” fluorescent channel) continuously increases, as cells transit to dead states. By comparing the relative intensity between NucGreen and NucBlue signals, semantic segmentation maps can be generated to label individual cell as either live or dead, as shown in Fig. 1e. The procedure of generating the semantic maps can be found in the *Supplemental Information Section 1*.

### Deep neural network architecture, training, validation and testing

With fluorescence-based semantic maps as ground truth, a deep neural network was trained to assign “live”, “dead”, or background labels to pixels in the input SLIM images. We employed a U-Net based on EfficientNet (E-U-Net) [30], with its architecture shown in Fig.2a. Compared to conventional U-Nets, the E-U-Net uses EfficientNet [30], a powerful network of relatively lower complexity, as the encoding part. This architecture allows for learning an efficient and accurate end-to-end segmentation model, while avoiding training a very complex network. The network was trained using a transfer learning strategy [31] with a finite training set. At first, the EfficientNet of E-U-Net (the encoding part) was pre-trained for image classification on a publicly available dataset ImageNet [35]. The entire E-U-Net was then further fine-tuned for a semantic segmentation task by using 899 labeled SLIM images as the training set and 199 labeled images as the validation set. The network training was performed by updating the weights of parameters in the E-U-Net using an Adam optimizer [36] to minimize a loss function that is computed in the training set. More details about the EfficientNet module and loss function can be found in the *Supplemental Information Section 2&3*. The network was trained for 100 epochs. At the end of each epoch, the loss function related to the being-trained network was evaluated, and the weights that yielded the lowest loss on the validation set were selected for the E-U-Net model. Figure 2d shows training and validation loss vs. number of epochs. The *Methods* section and Figs. 2a-c present more details about the E-U-Net architecture and network training.

**Figure 2.**
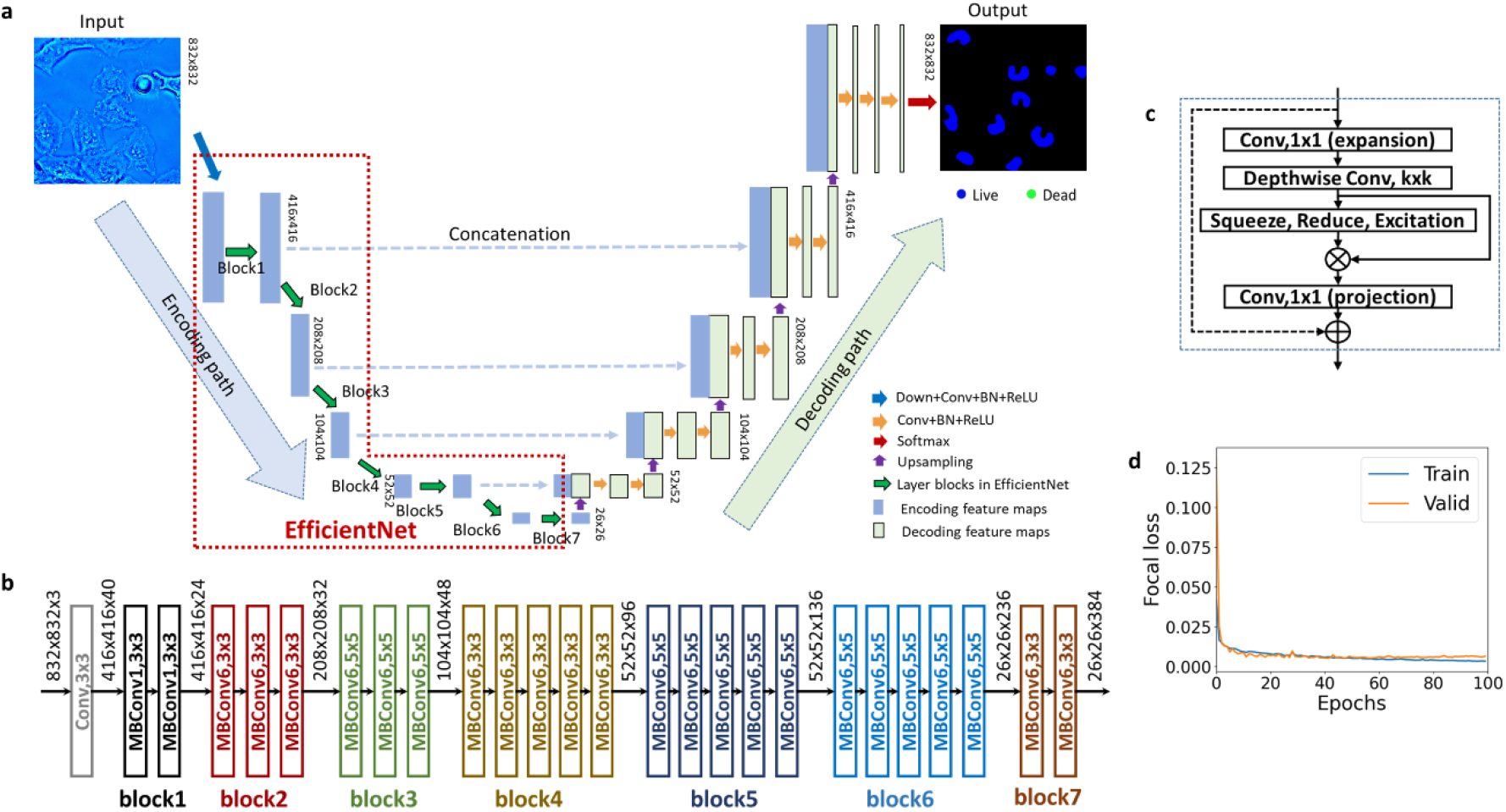
Principle of E-U-Net training. **a**. The E-U-Net. architecture includes an EfficientNet as the encoding path and five stages of decoding. The E-U-Net includes a Down+Conv+BN+ReLU block and 7 other blocks. The Down-Conv-BN-ReLU block represents a chain of down-sampling layer, convolutional layer, batch normalization layer, and ReLU layer. Similarly, the Conv+BN+ReLU is a chain of convolutional layer, batch normalization layer, and ReLU layer. **b** The network architecture of EfficientNet-B3. Different blocks are marked in different colors. They correspond to the layer blocks of EfficientNet in **a. c**. The major layers inside the MBConvX module. X =1 and X = 6 indicate the ReLU and ReLU6 are used in the module, respectively. The skip connection between the input and output of the module is not used in the first MBConvX module in each layer block. **d**. Training and validation loss vs epochs.

To demonstrate the performance of phase imaging with computational specificity (PICS) as a label-free live/dead assay, we applied the trained network to 200 SLIM images not used in training and validation. Figure 3a shows the three representative testing phase maps, whereas corresponding ground truth and PICS prediction are shown in Fig. 3b and Fig. 3c, respectively. This direct comparison indicates that PICS successfully identifies the region of cell nucleus with high confidence. As reported in previous publications, the conventional deep learning evaluation metrics focus on assessing pixel-wise segmentation accuracy, which overlooks some biologically relevant instances [37]. Here, we adopted an object-based evaluation metric, which relies on comparing the dominant semantic label between the predicted cell nuclei and the ground truth for individual nucleus. The confusion matrix and the corresponding evaluation (e.g. precision, recall and F1-score) are shown in Table. 1. A comparison with standard pixel-wise evaluation is included in Table. S2 in *Supplemental Information*. The entries of the confusion matrix are normalized with respect to the number of cells in each category. Using the average F1 score across all categories as an indicator of the overall performance, this PICS strategy reports a 96.7% confidence in distinguish individual live and dead HeLa cells.

**Table 1.** Evaluation of the E-U-Net performance. An object-based accuracy metric is used to estimate the deep learning prediction by comparing the dominant semantic label of HeLa cell nuclei with the ground truth. The entries of the confusion are normalized with respect to number of cells in each class.

| **Prediction \ Actual** | **Live (n= 1973)** | **Dead (n = 246)** |
| --- | --- | --- |
| **Live** | 98.8% | 2.4% |
| **Dead** | 1.2% | 97.6% |
| **Precision** | 99.6% | 91.2% |
| **Recall** | 98.8% | 97.6% |
| **F1 Score** | 99.1% | 94.3% |

**Figure 3.**
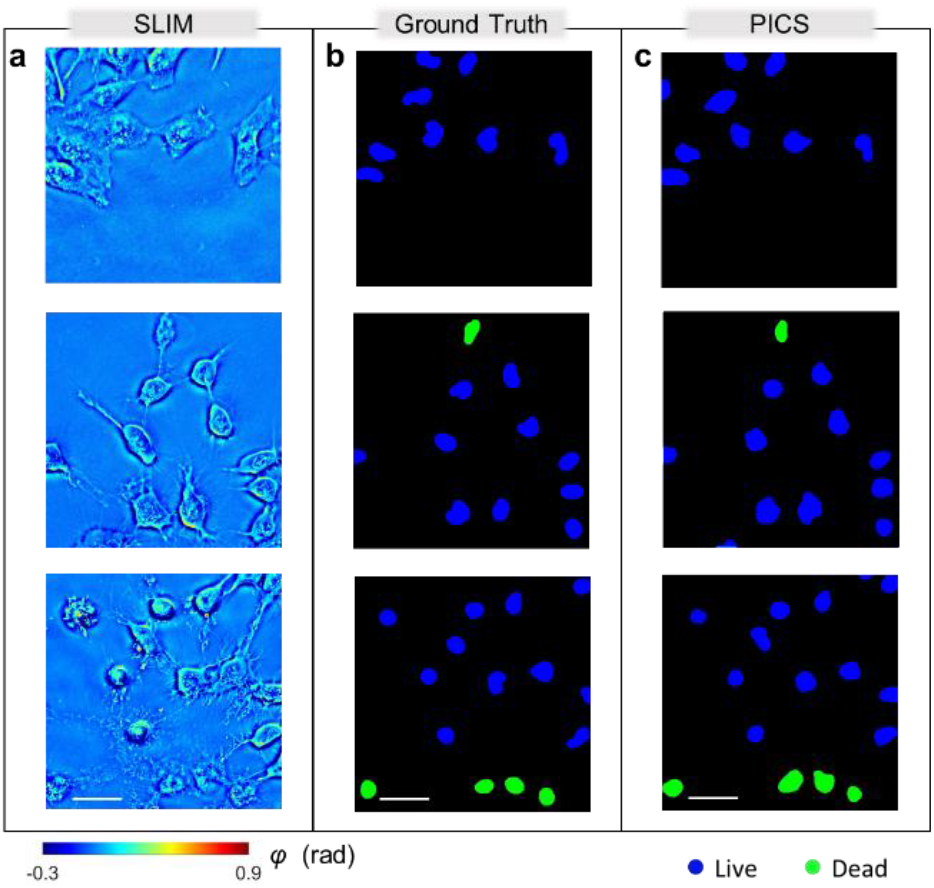
Results of E-U-Net on testing dataset. **a**. representative SLIM measurements of HeLa cells not used during training. **b**. The ground truth for viability of frames corresponding to **a. c**. The PICS prediction shows high level accuracy in segmenting the nuclear regions and inferring viability states. Scale bars: 50 microns

### PICS on CHO cells

Chinese hamster ovary (CHO) cells are often used for recombinant protein production, as it received U.S. FDA approval for bio-therapeutic protein production. Here, we demonstrate that our label-free viability assay approach is applicable to other cell lines of interest in pharmaceutical applications. CHO cells were plated on a glass bottom 6-well plate for optimal confluency. In addition to NucBlue/NucGreen staining, 1μM of staurosporine (apoptotic inducing reagent) solution was added to the culture medium. This potent reagent permeates cell membrane and disrupts protein kinase, cAMP, and lead to apoptosis in 4 -6 hours. The cells were then measured by SLIM and epi-fluorescence microscopy. To exclude the effect of necrosis, the cells were maintained in regular incubation condition throughout the experiment. After image acquisition, E-U-Net (EfficientNet-B7) training was immediately followed. In the training process, 1536 labeled SLIM images and 288 labeled SLIM images were used for network training and validation, respectively. The structure of EfficientNet-B7, training and validation loss can be found in Fig. S3a-b, respectively. The trained E-U-net was finally applied to 288 unseen testing images to test the performance of dead/viability assay. The procedure of imaging, ground truth generation, and training were consistent with the previous experiments.

Figure 4a shows the time-lapse SLIM image of CHO cells measured at t = 0, 2, and 10 hours after adding apoptosis reagent, and the corresponding viability map determined by fluorescence signal and PICS are plotted in Fig. 4b and Fig. 4c, respectively. In contrast to HeLa, the cell bodies became gradually fragmented during apoptosis. The visual comparison in Fig. 4 suggests that PICS yields good performance in extracting cell nucleus and predicting its viable state. Running an evaluation on individual cell, as shown in Table. 2, the network gives an average F-1 score of 94.9%. Furthermore, because most of the cells stay adherent, the PICS accuracy was not affected by cell confluence. In *Supplemental Information Section 4*, we show the evaluation and representative SLIM image of CHO cells under difference level of confluence.

**Table 2.** Evaluation of the E-U-Net performance on CHO with apoptosis reagents. The trained network yields high confidence in identifying live or apoptotic CHO cells. The entries of the confusion are normalized with respect to number of cells in each class.

| **Prediction \ Actual** | **Live (n= 2071)** | **Dead (n = 6328)** |
| --- | --- | --- |
| **Live** | 90.1% | 1.7% |
| **Dead** | 9.9% | 98.3% |
| **Precision** | 94.6% | 96.8% |
| **Recall** | 90.1% | 98.3% |
| **F1 Score** | 92.3% | 97.5% |

**Figure 4.**
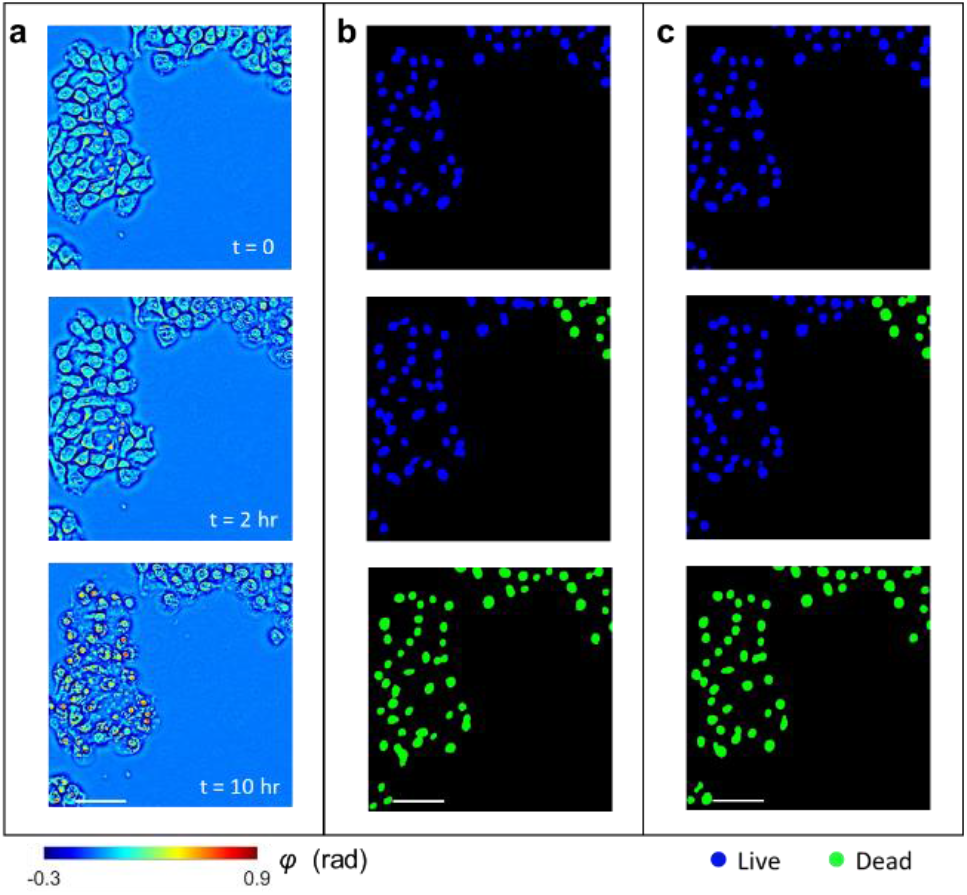
Results of PICS on adherent CHO cells. a. Time-lapse SLIM measurements of CHO cells measured at t= 0, 2, and 10 hours. The data was not used during training or validation. b. The ground truth for viability of frames corresponding to a. c. The PICS prediction shows high level accuracy in segmenting the nuclear regions and inferring viability states. Scale bars: 50 microns

### PICS on unlabeled HeLa cells

Performing viability assay on unlabeled cells essentially circumvents the cell injury effect caused by exogenous staining and produces an unbiased evaluation. To demonstrate this advantage, a fresh HeLa cell culture was prepared in a 6-well plate, transferred to the microscope stage, and maintained under room conditions. Half of the wells were mixed with viability assay reagents, where the viability was determined by both PICS and fluorescence imaging. The remaining wells did not contain reagents, such that the viability of these cells were only evaluated by PICS. The procedure of cell preparation, staining, and microscope settings were consistent with the previous experiments. We took measurements every 30 minutes, and the entire experiment lasted for 12 hours.

Figures 5a, c show SLIM images of HeLa cells with and without fluorescent reagents at t = 0, 2.5, and 12 hours, respectively, whereas the resulting PICS predictions are shown in Figs. 5b, d. *Supplemental Video 1* shows a time-lapse SLIM measurement, PICS prediction, and standard live-dead assay based on fluorescent measurements. *Supplemental Video 2* shows HeLa cells without reagents. As expected, the PICS method depicts the transition from live to dead state. In addition, the visual comparison from Figs. 5a-d suggests that HeLa cells with viability stains in the media appear smaller in size, and more rapidly entering the injured state, as compared to their label-free counterparts. Using TrackMate [38], an imageJ plugin, we were able to extract the trajectory of individual cells and track the cell morphology over time. As a result, the cell nucleus, area, and dry mass at each moment in time can be obtained by integrating the pixel value over the segmented area in the PICS prediction and SLIM image, respectively. We successfully tracked 57 labeled and 34 unlabeled HeLa cells. Figures 5e-f show the area and dry mass change (mean ± standard error), where the values are normalized with respect to the one at t=0. Running two sample t-tests, we found a significant difference in cell nuclear areas between the labelled and unlabeled cells, during the interval t=2 and t=7 hours (p < 0.05). Similarly, cell dry mass showed significant differences between the two groups during the time interval t=2 and t=5 hours (p < 0.05). As expected, these differences vanish as the cells in both groups eventually die.

**Figure 5.**
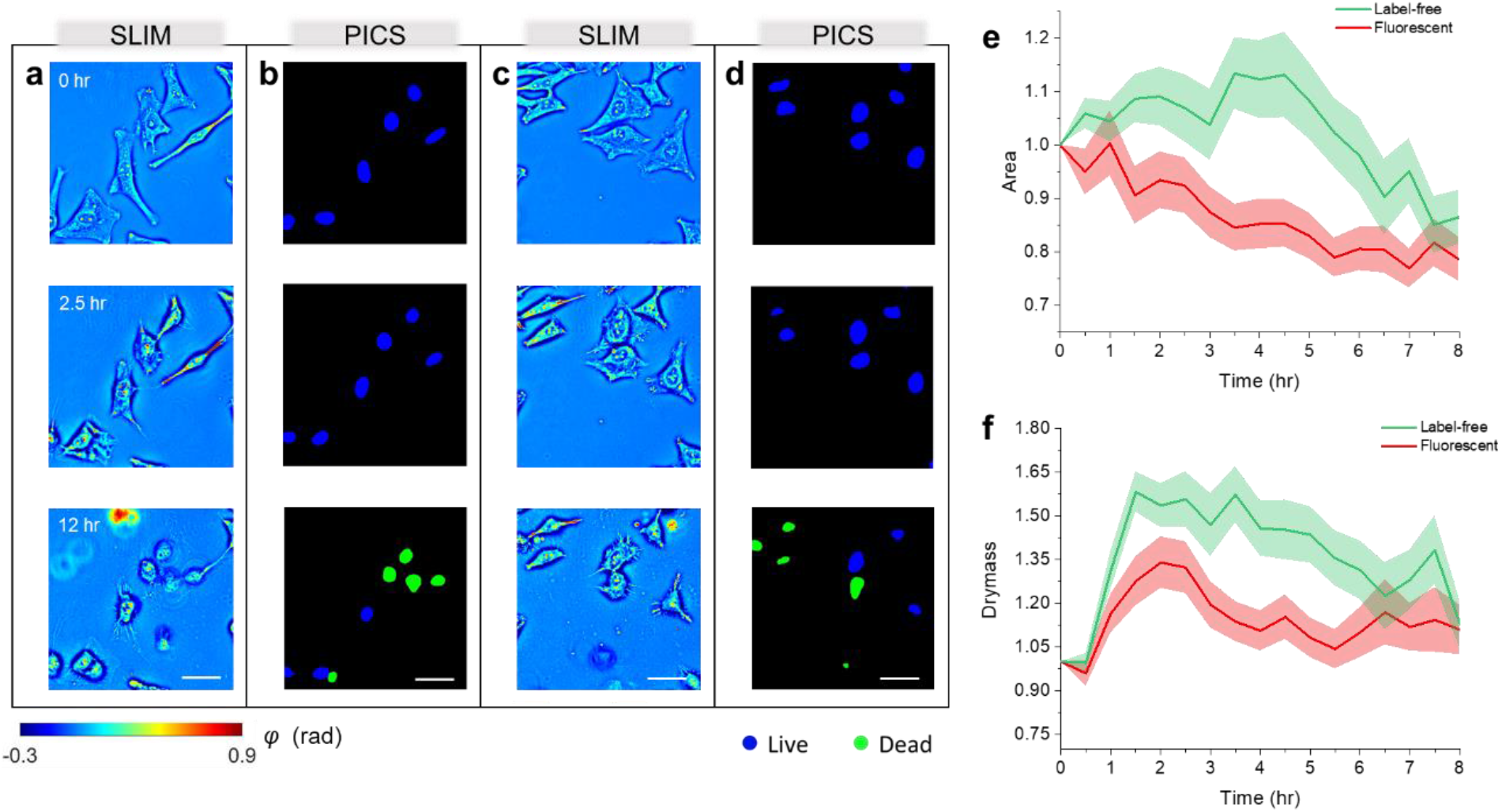
Viability of HeLa cells with and without reagent stains. **a**. SLIM images of cells recorded at 0, 2.5 and 12 hours after staining. **b**. The PICS prediction associated with the frames in **a. c**. SLIM images of unstained HeLa cells measured at same time points as **a. d**. The corresponding PICS prediction associated with the frames in **c. e**. Relative cell nuclear area change of tracked cells. The shaded region represents the standard error. **f**. Relative cell nucleus dry mass change. The shaded region represents of the standard error.

## Summary and Discussion

We demonstrated PICS as a method for high-throughput, label-free, unbiased viability assessment of adherent cells. This approach utilizes quantitative phase imaging to record high-resolution morphological structure of unstained cells, combined with deep learning techniques to extract intrinsic viability markers. Tested on HeLa and CHO adherent cultures, the proposed method reports 96.7% and 94.9% accuracy in segmenting the cell nucleus and classifying their viability state. By integrating the trained network on NVIDIA graphic processing units, the proposed label-free method enables real-time acquisition and viability prediction (see *Supplemental Video* 3 for a demonstration). One SLIM measurement and deep learning prediction takes ∼100 ms, which is approximately 8 times faster than the acquisition time required for fluorescence imaging with the same camera. Of course, the cell staining process itself takes time, approximately 15 minutes in our case. In addition, results suggest that PICS rules out the adverse effect on cell function caused by the exogenous staining, which is potentially beneficial for the unbiased assessment of cellular activity over long periods of time (*e*.*g*., many days). Of course, this approach can be applied to other cell types and cell death mechanisms.

Label-free imaging methods are valuable to study biological samples without destructive fixation or staining. The capabilities of PICS can be furthermore enriched by combining with other imaging modalities. For example, by employing infrared spectroscopy, the bond-selective transient phase imaging measures molecular information associated with lipid droplet and nucleic acids [39]. In addition, harmonic optical tomography can be integrated into an existing QPI system to report both non-centrosymmetric structures [40]. These additional chemical signals would potentially enhance the effective learning and produce more biophysical information. We anticipate that the PICS method will benefit high-throughput cell screening for a variety of applications, ranging from basic research to therapeutic development and protein production in cell reactors [8].

## Methods

### HeLa cell preparation

HeLa cervical cancer cells (ATCC <u>CCL-</u>2™) and Chinese hamster ovary (CHO-K1 ATCC <u>CCL-61</u> ™) cells were purchased from ATCC and kept frozen in liquid nitrogen. Prior to the experiments, we thawed and cultured the cells into T75 flask in Dulbecco’s Modified Eagle Medium (DMEM with low glucose) containing 10% fetal bovine serum (FBS) and incubated in 37°C with 5% CO2. As the cells reach 70% confluence, the flask was washed thoroughly with phosphate-buffered saline (PBS) and trypsinized with 3 mL of 0.25% (w/v) Trypsin EDTA for three minutes. When the cell starts to detach, the cells were suspended in 5 mL DMEM and passaged onto a glass bottom 6 well plate to grow. For the purpose of the study, CHO cells were plated in three different confluency levels: high (60000 cells), medium (30000 cells) and low (15000 cells). HeLa and CHO cells were then imaged after two days.

### SLIM imaging

The SLIM optical setup in shown in Fig. 1a. Detailed design and calibration procedure have been described previously [28]. In brief, the microscope is built upon an inverted phase contrast microscope using a SLIM module (CellVista SLIM Pro; Phi Optics) attached to the output port. Inside the module, a spatial light modulator (Meadowlark Optics) is placed at the system pupil plane via a Fourier transform lens to constantly modulate the phase delay between the scattered and incident light. By recording four intensity images with phase shifts of 0, *π*/2, π, and 3*π*/2, a quantitative phase map, *φ*, can be computed by combining the 4 acquired frames in real-time.

For both SLIM and fluorescence imaging, cultured cells were measured by a 40× objective, and the images were recorded by a CMOS camera (ORCA-Flash 4.0; Hamamatsu) with a pixel size of 6.5 μm. In this experiment, acquisition time of each SLIM and fluorescent measurement is 50 ms and 400 ms, respectively. For deep learning training and predicting, the recorded SLIM images were downsampled by a factor of 2. This step saves computational cost and does not sacrifice information content.

### E-U-Net architecture and network training

The E-U-Net is a U-Net-like fully convolutional neural network that performs an efficient end-to-end mapping from SLIM images to the corresponding probability maps, from which the desired segmentation maps are determined by use of a softmax decision rule. Different from conventional U-Nets, the E-U-Net uses a more efficient network architecture, EfficientNet [30], for feature extraction in the encoding path. Here, EfficientNets refers to a family of deep convolutional neural networks that possess a powerful capacity of feature extraction but require much fewer network parameters compared to other state-of-the-art network architectures, such VGG-Net, ResNet, Mask R-CNN, etc. The EfficientNet family includes eight network architectures, EfficientNet-B0 to EfficientNetB7, with an increasing network complexity. EfficientNet-B3 and EfficientNet-B7 were selected for training E-U-Net on HeLa cell images and CHO cell images, respectively, considering they yields the most accurate segmentation performance on the validation set among all the eight EfficientNets. See Supplemental information and Fig.2b-2c for more details about the EfficientNet-B3 and EfficientNet-B7.

The E-U-Net was trained with randomly cropped patches of 512×512 pixels drawn from the training set by minimizing a loss function with an Adam optimizer [36]. In regard to Adam optimizer, the exponential decay rates for 1^st^ and 2^nd^ moment estimates were set to 0.9 and 0.999, respectively; a small constant *ε* for numerical stability was set to 10^−7^. See *Supplemental information* for more details related to the loss function. The batch size was set to 14 and 4 for training E-U-net on the HeLa cell images and CHO cell images, respectively. The learning rate was initially set to 5×10^−4^. At the end of each epoch, the loss of the being-trained E-U-Net was computed on the whole validation set. When the validation loss did not decrease for 10 training epochs, the learning rate was multiplied by a factor of 0.8. This validation loss-aware learning rate decaying strategy benefits for mitigating the overfitting issue that commonly occurs in deep neural network training. Furthermore, data augmentation techniques, such as random cropping, flipping, shifting, and random noise and brightness adding etc., were employed to augment training samples on-the-fly for further reducing the overfitting risk. The E-U-Net was trained for 100 epochs. The parameter weights that yield the lowest validation loss were selected, and subsequently used for model testing and further model investigation.

The E-U-Net was implemented using the Python programming language with libraries including Python 3.6 and Tensorflow 1.14. The model training, validation and testing were performed on a NVIDIA Tesla V100 GPU of 32 GB VRAM.

## Data availability

The images and generated that support the findings of this study are available from the corresponding author upon request.

## Code availability

The MATLAB script for semantic segmentation maps and evaluation are available from the corresponding author upon request. The trained E-U-Net models along with sample testing images are available for download at https://github.com/shenghh2020/label-free-viability-assay.

## Acknowledgement

This work is sponsored in part by the National Science Foundation (0939511, 1353368), and the National Institute of Health (R01CA238191, R01GM129709). C. H. is supported by National Institute of Health – Tissue Microenvironment (TiMe) training program (T32EB019944).

## Author contribution

C. H. designed the experiment, collected data, generated semantic segmentation maps, performed deep learning testing, and conducted data analysis. S. H. led the deep learning training and validation. Y. L. prepared cell cultures. Y. H. performed software integration. E. M. K prepared the script for PICS evaluation. C. H. and S. H wrote the manuscript with input from all authors. M. A. A. supervised the deep learning. G. P. supervised the project.

## List of supplemental materials

1. Supplemental Information
2. Supplemental Movie 1: Animation of HeLa cells with viability reagents measured by SLIM (left), corresponding PICS prediction (middle), and standard viability assay (right)
3. Supplemental Movie 2: Animation of HeLa cells without staining reagents measured by SLIM (left), and corresponding PICS prediction (right)
4. Supplemental Movie 3: Real-time SLIM measurement and PICS viability prediction

