## Supplemental_Information for "Live-dead assay on unlabeled cells using phase imaging with computational specificity"

### 1. Semantic map generation

Semantic segmentation maps were generated in MATLAB with a customized script. First, for each NucBlue and NucGreen image pair, an adaptive thresholding was applied to separate the cell nucleus and background, where the segmented cell nuclei were obtained by computing the union of the binarized fluorescent image pair. We removed the segmentation artifacts by filtering out the tiny objects below the size of a typical nucleus. Next, using on the segmentation masks, we calculated the ratio between the NucGreen and NucBlue fluorescence signal. A histogram of the average ratio within the cell nucleus is plotted in Fig. S1, where three distinctive peaks were observed corresponding to the live, injured and dead cells. Because NucGreen/NucBlue reagent is only designed for live and dead classification, the histogram of injured cells is partially overlapped with the live cells. By selecting a threshold value that gives the lowest histogram count between dead and injured cells, we assigned label "live" to all live and injured cells, while the remaining cells as "dead".

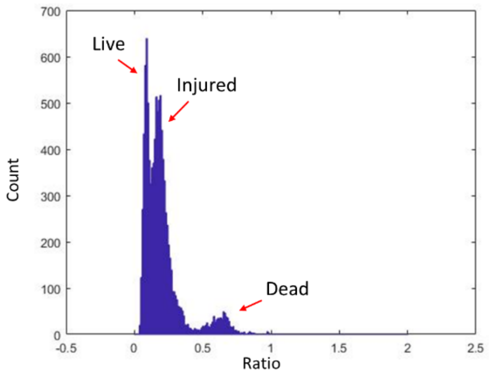

**Figure S1.** Histogram of fluorescence signal ratio.

### 2. EfficientNet

The MBConvX is the principle module in an EfficientNet. It approximately factorizes a standard convolutional layer into a sequence of separable layers to shrink the number of parameters needed in a convolution operation while maintaining a comparable ability of feature extraction. The separable layers in a MBConvX module are shown in Fig. 2c. Here, MBConv1(X=1) and MBConv6 (X=6) indicate that a ReLU layer and ReLU6 layer are employed in this module, respectively. ReLU6 is a modification of the rectified linear unit, where the activation is limited to a maximum size of 6. A MBConvX module in Fig. 2b may include a down-sampling layer, which can be inferred by the indicated feature map dimensions. The first MBConvX in each layer block does not contain a skip connection between its input and output (indicated as a dash line in Fig. 2c), since the input and output of that module have different sizes.

### 3. Loss function

Given a set of $B$ training images of $M \times N$ pixels and their corresponding ground truth semantic segmentation maps, the E-U-Net is trained by minimizing a loss function that is computed between the predicted probability maps and the corresponding ground truth label maps in the training set. The loss function is defined as the combination of focal loss [1] and dice loss [2]:

$$L_{Focal\_loss} = -\frac{1}{B}\sum_{i=1}^{B}\frac{1}{MN}\sum_{x\in\Omega}\left[1-y_i(x)^T p_i(x)\right]^{\gamma} y_i(x)^T \log_2 p_i(x), \quad [1]$$

$$L_{Dice\_loss} = 1-\frac{1}{3}\sum_{c=0}^{2}\frac{2TP_c}{2TP+FP_c+FN_c} \quad [2]$$

$$L_{combined} = \alpha L_{Focal\_loss} + \beta L_{Dice\_loss} \quad [3]$$

In the focal loss $L_{Focal\_loss}$, $\Omega = \{(1,1),(1,2),...,(M,N)\}$ is the set of spatial locations of all the pixels in a label map. $y_i(x) \in \{[1,0,0]^T, [0,1,0]^T, [0,0,1]^T\}$ represents the ground truth label of the pixel *x* related to the $i^{th}$ training sample, and the three one-hot vectors correspond to the live, dead and, background classes, respectively. Accordingly, the probability vector $\mathbf{P}_i(x) \in \mathbb{R}^3$ represents the corresponding predicted probabilities of belonging the three classes. $\left[1 - y_i(x)^T p_i(x)\right]^\gamma$ is a classification error-related weight that reduces the relative cross entropy $y_i(x)^T \log_2 p_i(x)$ for well-classified pixels, putting more focus on hard, misclassified pixels. In this study, $\gamma$ was set to be the default value of 2 as suggested in Ref. [1]. As the dice loss $L_{Dice\_loss}$, the $TP_c$, $FP_c$, and $FN_c$ are the number of true positives, that of false positives, and that of false negatives, respectively, related to all pixels of viability class $c \in \{0,1,2\}$ in the B images. Here, $c$ = 0, 1, and 2 correspond to the live, dead and background classes, respectively. In the combined loss function, $\alpha, \beta \in \{0,1\}$ are two indicators that controls whether to use focal loss and dice loss in the training process, respectively. In this study, $\alpha, \beta$ was set to [1, 0] and [1, 1] for training the E-U-Net on HeLa cell dataset and CHO cell dataset, respectively. The choices of [$\alpha, \beta$] were determined by segmentation performance of the trained E-U-Net on the validation set. The combined loss is minimized with an Adam optimizer.

### 4. PICS Performance on adherent CHO cells under different confluence

As discussed in the manuscript, live CHO cell culture was prepared in a 6-well plate at three confluence levels, staurosporine solution was added into the culture medium to introduce apoptosis. Figure. S2 show SLIM image of high, intermediate, and low confluence CHO cells measured at t = 0. Although, aggregating into clusters, the cell shape and boundary can be easily identified. All SLIM images were combined for training and validation. In testing, we estimated the PICS performance vs. cell confluence, and the results are summarized in Table. S1a-c.

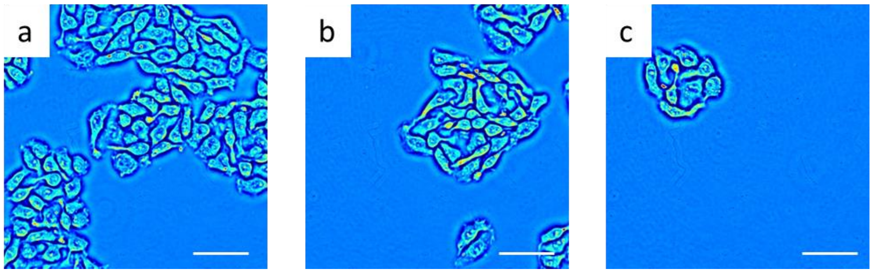

**Figure S2**. SLIM images of high (a), intermediate (b), and low (c) confluence CHO cells. Scale bar: 50 µm in space

a

| High | Live | Dead |
|---|---|---|
| **Precision** | 90.5% | 98.5% |
| **Recall** | 94.7% | 97.2% |
| **F1 Score** | 92.6% | 97.8% |

b

| Intermediate | Live | Dead |
|---|---|---|
| **Precision** | 87.5% | 98.7% |
| **Recall** | 96.3% | 95.2% |
| **F1 Score** | 91.7% | 96.9% |

c

| Low | Live | Dead |
|---|---|---|
| **Precision** | 96.2% | 97.1% |
| **Recall** | 90.1% | 98.9% |
| **F1 Score** | 93.1% | 97.9% |

**Table S1.** PICS performance vs. CHO cell confluence

Figure S3. **CHO cells viability training with EfficientNet-B7.** (a) The network architecture of EfficientNet-B7. (b) Training and validation focal losses vs number of epochs.

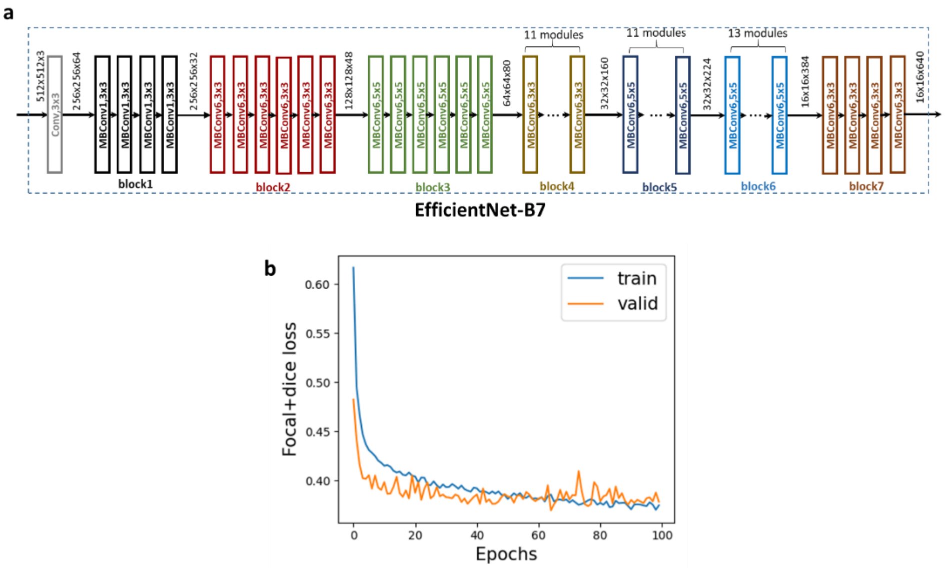

Table S2. **Pixel-wise evaluation of the trained E-U-Net.** The confusion matrix and F1 score of the E-U-Net performance evaluated on pixels in testing images. Due to the fact that the E-U-Net prediction assigns multiple labels to one cell nucleus, we converted the pixel-wise classification into cell-wise classification, which is more relevant biologically (Table 1 in the main text).

| Prediction \ Actual | Live | Dead | Background |
|---|---|---|---|
| Live | 3699117 | 2591 | 1068559 |
| Dead | 1422 | 221396 | 187715 |
| Background | 696478 | 44028 | 62668870 |
| **F1 Score** | 80.7% | 65.3% | 98.4% |
